# Wnt5a-Ror2 signaling activates p62-Nrf2 axis in reactive astrocytes after brain injury

**DOI:** 10.1101/2023.04.19.537450

**Authors:** Mitsuharu Endo, Yuki Tanaka, Mayo Fukuoka, Hayata Suzuki, Yasuhiro Minami

## Abstract

In the brains under pathological conditions, astrocytes become reactive astrocytes that exhibit various context-dependent functions through the regulation of specific signaling pathways and transcriptional mechanisms in response to environmental changes. Reactive astrocytes induced in injured brains begin proliferating and play a role in promoting protection and repair of damaged tissues, but the relationship between the proliferative characteristics and tissue-protective and repair functions of reactive astrocytes remains unclear. Here, we show that growth factor signaling elicited by bFGF and HB-EGF, whose expression is up-regulated in the injured brains, acts synergistically with inflammatory cytokine signaling in astrocytes, thereby markedly up-regulating gene expression of the Ror-family protein Ror2, a receptor for Wnt5a. Activation of Wnt5a-Ror2 signaling in astrocytes results in intracellular accumulation of phosphorylated p62, thereby activating antioxidative transcription factor Nrf2. Finally, we provide evidence demonstrating that forced activation of Wnt5a-Ror2-p62-Nrf2 signaling axis in astrocytes reduces cellular damage caused by hemin, a degradation product of hemoglobin, and promotes repair of the damaged blood brain barrier after brain hemorrhage.

## Introduction

Astrocytes are a type of glial cells in the central nervous system and play important roles in maintaining brain homeostasis. Under pathological conditions, they become reactive astrocytes characterized by cellular hypertrophy and increased expression of various genes, including glial fibrillary acidic protein (GFAP). Recent studies have revealed that reactive astrocytes exhibit heterogeneity in their gene expression profiles, performing a variety of functions depending on the pathological context (Zamanian *et al*, 2012; Liddelow *et al*, 2017). In the injured central nervous system, including traumatic brain injury (TBI), stroke, and intracerebral hemorrhage (ICH), reactive astrocytes have been shown to initiate proliferation and contribute to tissue protection and repair (Morizawa *et al*, 2017; Chiareli *et al*, 2021; Williamson *et al*, 2021). Although reactive astrocytes do not necessarily exhibit proliferative capacity, proliferation appears to be a key feature of reactive astrocytes induced in the injured brains. Ganciclovir-mediated depletion of proliferating reactive astrocytes in the injured brains increases the spread of inflammatory cells from damaged areas into neighboring unaffected tissues (Myer *et al*, 2006). Proliferative reactive astrocytes have also been shown to promote repair of damaged blood vessels and blood brain barrier (BBB) (Williamson *et al*., 2021). However, the relationship between proliferative properties and tissue-protective functions of reactive astrocytes during tissue repair remains unclear.

The Ror-family receptor Ror2 acts as a receptor for Wnt5a and plays crucial roles in the generation of various tissues during development (Minami *et al*, 2010; Endo *et al*, 2022). In the developing neocortices, Ror2 is expressed highly in neural stem/progenitor cells and mediates the regulation of their stemness, including proliferative ability (Endo *et al*, 2012). In adult brains, expression of *Ror2* is maintained at lower levels, but is up-regulated in reactive astrocytes following brain injury, and Ror2 signaling promotes their proliferation (Endo *et al*, 2017). We have shown that expression of several growth factors, including bFGF, is up-regulated in the injured brains (Endo *et al*., 2017), and that bFGF stimulation can induce expression of *Ror2* in fibroblasts via activation of E2F1 (Endo *et al*, 2020). Although there are several findings on the transcriptional mechanism of *Ror2*, the detailed regulatory mechanisms in reactive astrocytes are not fully understood, and it is also largely unknown about how downstream signaling pathways mediated by Ror2, whose expression is induced by transcriptional activation, can contribute to tissue protection and repair through the regulation of astrocyte functions under oxidative conditions following brain injury with hemorrhage.

Here, we show that Ror2-expressing proliferative reactive astrocytes are located in the vicinity of inflammatory cells, and that growth factor signaling, including bFGF and HB-EGF signaling, and inflammatory cytokine signaling, including IL-1β and TNF-α signaling, act cooperatively to engage E2F1-mediated transcription of *Ror2* in cultured astrocytes. Furthermore, we found that Wnt5a-Ror2 signaling induces activation of p62-Nrf2 signaling axis in astrocytes following stimulation by inflammatory cytokines together with bFGF. Phosphorylation of p62 at Ser-351 has been shown to result in activation of Nrf2, an antioxidative transcription factor, through its binding to Keap1 (Ichimura *et al*, 2013). With this respect, we provide evidence showing that p62 is conventionally degraded rapidly by autophagy in astrocytes, but activation of Wnt5a-Ror2 signaling suppresses autophagic degradation of p62, thereby promoting intracellular accumulation of phosphorylated p62. Intriguingly, analyses of a mouse ICH model with collagenase injection further exemplified that Wnt5a-Ror2 signaling indeed induces Nrf2 activation mediated by phosphorylated p62 in reactive astrocytes surrounding injured area, presumably contributing to repair of the damaged BBB.

## Results

### Ror2-expressing proliferative reactive astrocytes are adjacent to inflammatory immune cells in the injured brains

We have previously shown that bFGF signaling is activated in reactive astrocytes in the mouse brains after stab wound (SW) injury, and that expression of *Ror2* is up-regulated in these proliferative reactive astrocytes (Endo *et al*., 2017). By analyzing a resource of reactive astrocyte transcriptome (GEO: GSE35338) (Zamanian *et al*., 2012), we further obtained evidence that expression levels of *Ror2* are up-regulated significantly in proliferative reactive astrocytes following ischemic brain injury (middle cerebral artery occlusion; MCAO), but marginally in non-proliferative reactive astrocytes following LPS-induced brain inflammation (Appendix Fig. S1), suggesting that up-regulation of *Ror2* might play an important role in regulating functions of reactive astrocytes associated with tissue repair.

To reveal the role of Ror2-expressing proliferative reactive astrocytes in the injured brains, we first determined their location in the mouse brains with SW injury by immunohistochemical analysis. We found that Ror2-expressing proliferative reactive astrocytes are located close to Iba1-expressing immune cells, including microglias and macrophages, accumulated at the lesion core (Fig. 1A-C). It has been shown that activated microglias/macrophages secrete inflammatory cytokines, including IL-1 and TNF, and thereby affect gene expression profiles of reactive astrocytes. Indeed, expression levels of *IL-1β* and *TNF-α* were increased dramatically in the brains within 1 day after injury (Fig. 1D). Expression analyses of FACS-sorted cells from the intact and injured brains revealed that *IL-1β* and *TNF-α* are expressed predominantly in activated macrophages (CD45^high^) and microglias (CD45^low^), respectively (Fig. 1E and Appendix Fig. S2). We further examined which cell types serve as sources of growth factors, and found that expression levels of *bFGF* and *HB-EGF* are up-regulated in the reactive astrocytes themselves rather than immune cells and oligodendrocytes (Fig. 1E and Appendix Fig. S2). These results raise a possibility that function of Ror2-expressing proliferative reactive astrocytes might be affected by inflammatory cytokines together with growth factors.

**Figure 1.**
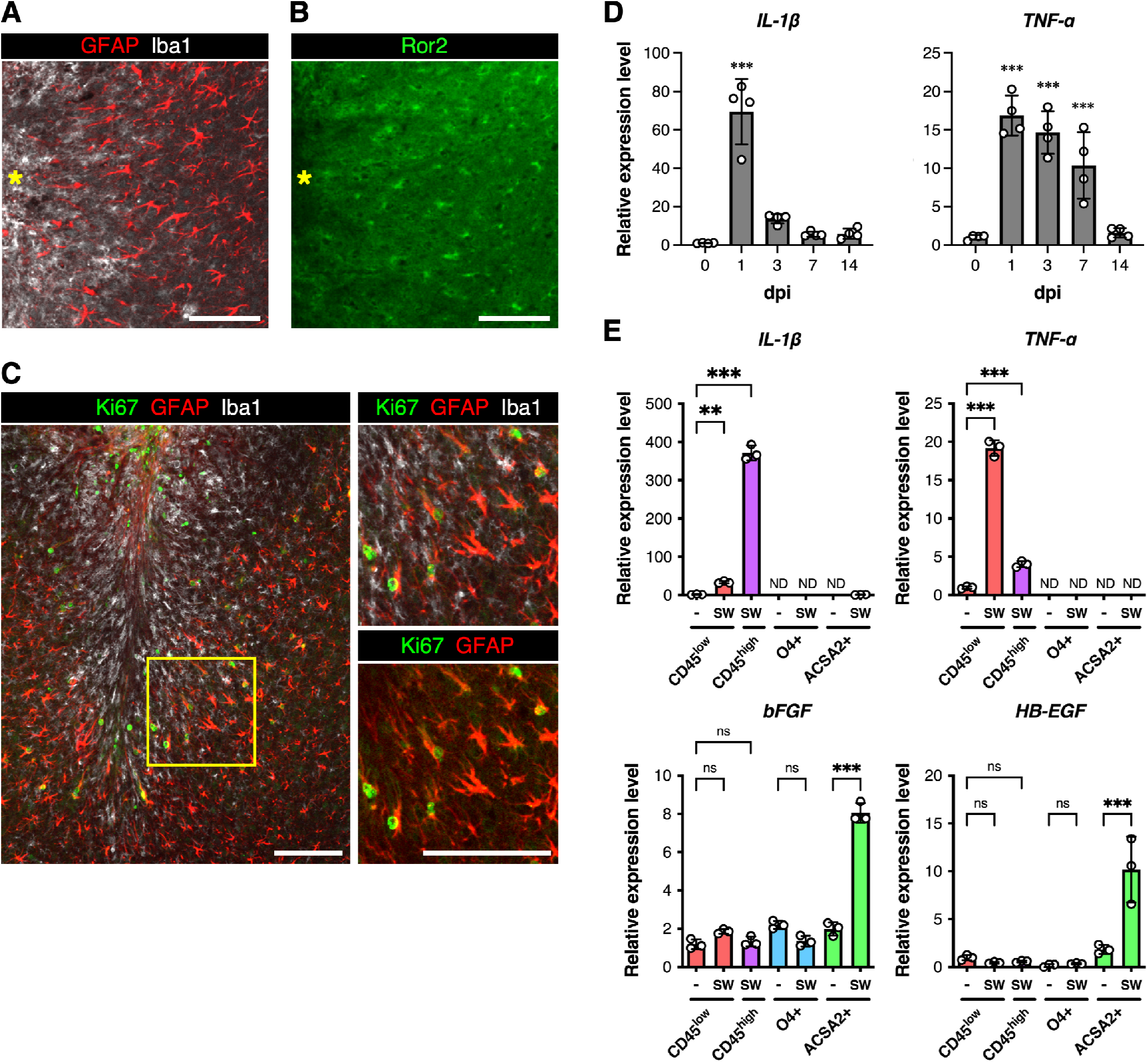
Ror2-expressing proliferative reactive astrocytes surround inflammatory cells in the injured brains. (A), (B) Neocortical tissues on day 5 after SW injury were stained with antibodies against GFAP (red, A), Iba1 (white, A), and Ror2 (green, B). Asterisks (yellow) indicate the injury site. Scale bars: 100 μm. (C) Neocortical tissues on day 5 after SW injury were stained with antibodies against Ki67 (green), GFAP (red), and Iba1 (white). The right panels are magnified views of the square in the left panel. Scale bars: 100 μm. (D) qRT-PCR analyses of *IL-1β* and *TNF-α* in injured [1, 3, 7, 14 days post-injury (dpi)] and uninjured (0 dpi) tissues of mouse neocortices. Data are shown as the mean ± SD (n = 4 animals per group). \*\*\**P*<0.001 (Dunnett’s test). (E) qRT-PCR analyses of *IL-1β*, *TNF-α*, *bFGF*, and *HB-EGF* in microglias (CD45^low^, red bars), macrophages (CD45^high^, purple bars), oligodendrocytes (O4+, blue bars), and astrocytes (ACSA2+, green bars) isolated from injured (SW) and uninjured (-) tissues of mouse neocortices. Data are shown as the mean ± SD (n = 3 animals per group). ND: not detectable. \*\**P*<0.01, \*\*\**P*<0.001 (Tukey’s test), ns: not significant.

### Expression of *Ror2* is induced by E2F1 through the synergistic action of growth factors and inflammatory cytokines

Next, we examined the possible involvement of IL-1β and TNF-α in regulating expression of *Ror2* by using cultured astrocytes. We found that mRNA and protein levels of Ror2 can be increased by stimulation with IL-1β and/or TNF-α, and further enhanced by co-stimulation with bFGF or HB-EGF (Fig. 2A-C), while expression levels of its putative ligand Wnt5a were unaffected (Fig. 2C). We have reported that E2F1 can mediate bFGF-induced expression of *Ror2* by binding to its promoter in NIH/3T3 fibroblasts (Endo *et al*., 2020). To investigate whether E2F1 also mediates expression of *Ror2* in astrocytes stimulated with IL-1β and TNF-α (hereafter referred to as I/T when both are added simultaneously), we expressed a dominant negative mutant of E2F1 fused with estrogen receptor (ERT2-E2F1-1′C). Up-regulation of *Ror2* induced by stimulation with I/T in the absence or presence of bFGF was significantly inhibited after nuclear translocation of ERT2-E2F1-1′C by an addition of 4-hydroxytamoxifen (OHT) (Fig. 2D). Similar results were obtained by suppressing expression of *Ep400*, encoding a critical component of the histone acetyltransferase complex involved in E2F-mediated transcriptional activation (Fig. 2E) (Taubert *et al*, 2004). Indeed, expression levels of *E2F1* and amounts of E2F1 bound to the *Ror2* promoter were increased in astrocytes stimulated with I/T together with bFGF (Figs. 2F and EV1). Interestingly, expression level of *CyclinE1*, a representative cell cycle-related target gene of E2F1, was not affected by stimulation with I/T in bFGF-treated astrocytes (Fig. EV1). Furthermore, up-regulated expression of *Ki67* by bFGF was rather reduced when stimulated with I/T (Fig. EV1), indicating that inflammatory cytokines can prevent bFGF-induced cell cycle progression in cultured astrocytes. These results suggest that growth factors and inflammatory cytokines act synergistically to up-regulate expression of *Ror2* through the binding of E2F1 to the *Ror2* promoter in astrocytes irrespective of their cell cycle progression (Fig. 2G).

**Figure 2.**
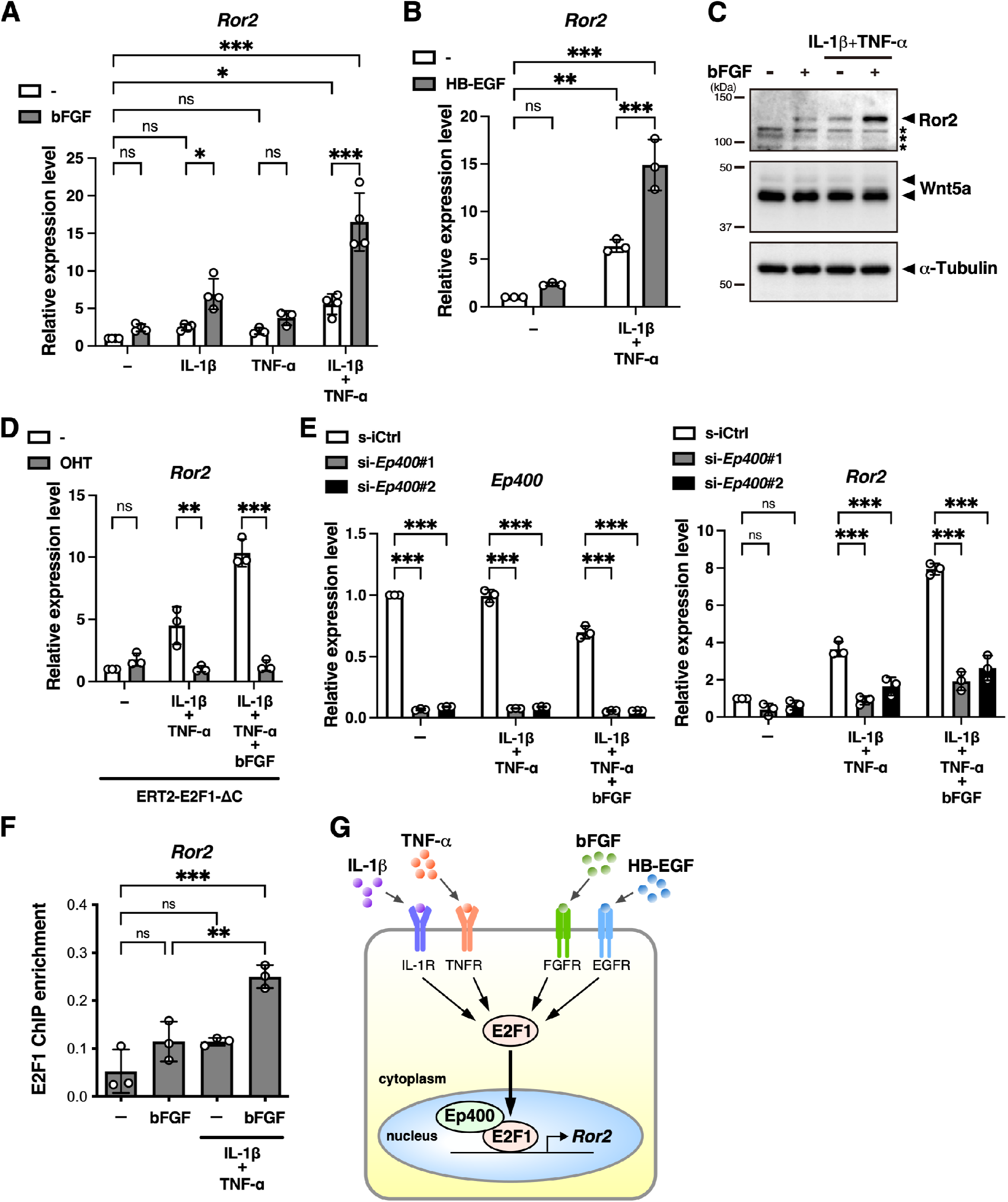
Expression of Ror2 is induced by E2F1 through the synergistic action of growth factors and inflammatory cytokines. (A-C) Differentiated astrocytes were stimulated with IL-1β, TNF-α or both in the presence or absence of bFGF (A) or HB-EGF (B) for 3 days. (A, B) Expression levels of *Ror2* were analyzed by qRT-PCR. Relative expression values were determined by defining the expression level of *Ror2* in untreated cells (-) [in the absence of bFGF (A) or HB-EGF (B)] as 1.0. The values are the mean ± SD of three to four independent experiments. \**P*<0.05, \*\**P*<0.01, \*\*\**P*<0.001 (Tukey’s test), ns: not significant. (C) Expression levels of Ror2, Wnt5a, and α-Tubulin were analyzed by Western blotting. Asterisks indicate non-specific bands. (D) Differentiated astrocytes expressing ERT2-E2F1-ΔC were treated with 300 nM 4-hydroxytamoxifen (OHT) for 2 h or left untreated (-), and then stimulated with IL-1β and TNF-α in the presence or absence of bFGF for 3 days. (E) Differentiated astrocytes transfected with *Ep400* siRNAs (si-*Ep400*#1 and si-*Ep400*#2) or control siRNA (si-Ctrl) were stimulated with IL-1β and TNF-α in the presence or absence of bFGF for 3 days. (D, E) Expression levels of *Ror2* were analyzed by qRT-PCR. Relative expression values were determined by defining the expression level of *Ror2* in each control group as 1.0. The values are the mean ± SD of three independent experiments. \*\**P*<0.01, \*\*\**P*<0.001 (Tukey’s test), ns: not significant. (F) Cross-linked chromatins were prepared from untreated cells (-) or cells stimulated with bFGF in the presence or absence of IL-1β and TNF-α for 3 days. The binding of E2F1 on the *Ror2* promoter was analyzed by ChIP analysis. The values are the mean ± SD of three independent experiments. \*\**P*<0.01, \*\*\**P*<0.001 (Tukey’s test), ns: not significant. (G) A schematic representation of E2F1-mediated transcriptional activation of *Ror2* in astrocytes stimulated with inflammatory cytokines and growth factors.

### Wnt5a-Ror2 signaling promotes nuclear accumulation of Nrf2

To determine the function of Ror2-mediated signaling in reactive astrocytes in injured brains, we explored genes induced by IL-1β together with bFGF in a manner depending on the expression of *Ror2* in cultured astrocytes by RNA-Seq analysis and compared those with genes up-regulated in reactive astrocytes in the injured brains. We identified 56 genes whose expression was up-regulated by co-stimulation with IL-1β and bFGF that were differentially down-regulated in si-*Ror2*-treated cells compared to si-Ctrl-treated ones (Table EV1). Among them, 13 genes were up-regulated significantly in reactive astrocytes induced by MCAO, but not in those induced by LPS administration (Table EV1). In particular, *heme oxygenase 1* (*Hmox1/HO-1*) was up-regulated highly in reactive astrocytes induced by SW brain injury (Table EV1), and increased expression of HO-1 was detected in Ror2-expressing proliferative reactive astrocytes surrounding the lesion core (Fig. EV2). By qRT-PCR analyses, we confirmed that expression levels of *HO-1* were increased in cultured astrocytes stimulated with I/T, and enhanced drastically by co-stimulation with bFGF (Fig. 3A), and that those were reduced by suppressed expression of *Ror2* or *Ep400*, but not *Ror1*, a gene encoding another member of the Ror-family receptors (Fig. 3B and Appendix Fig. S3A-C). Although expression level of Wnt5a was unaffected in cultured astrocytes after stimulation with IL-1β, TNF-α and bFGF (hereafter referred to as I/T/F) (Fig. 2C), expression levels of *HO-1* were decreased or increased by *Wnt5a* knockdown or recombinant Wnt5a treatment, respectively (Fig. 3B and C and Appendix Fig. S3D).

**Figure 3.**
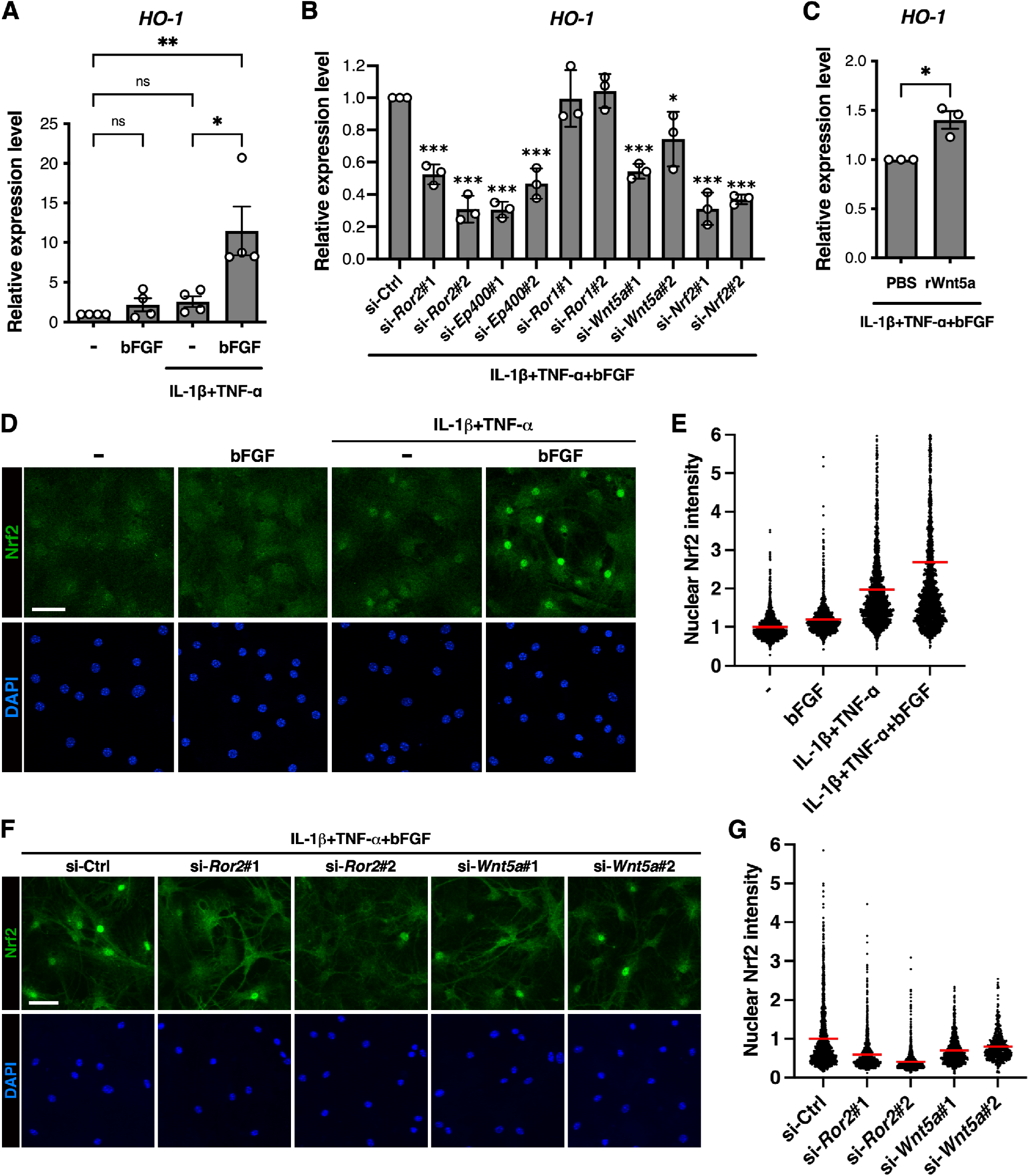
Wnt5a-Ror2 signaling promotes nuclear accumulation of Nrf2. (A) Differentiated astrocytes were stimulated with bFGF in the presence or absence of IL-1β and TNF-α for 3 days. (B) Differentiated astrocytes transfected with the indicated siRNAs were stimulated with IL-1β, TNF-α, and bFGF for 3 days. (C) Differentiated astrocytes were stimulated with IL-1β, TNF-α, and bFGF together with either recombinant Wnt5a (rWnt5a) (200 ng/mL) or its vehicle (PBS) alone for 3 days. (A-C) Expression levels of *HO-1* were analyzed by qRT-PCR. Relative expression values were determined by defining the expression levels of *HO-1* in each control group as 1.0. The values are the mean ± SD of three independent experiments. \**P*<0.05, \*\**P*<0.01, \*\*\**P*<0.001 (Tukey’s test for A; Dunnett’s test for B; Student’s *t*-test for C), ns: not significant. (D, E) Differentiated astrocytes were stimulated with bFGF in the presence or absence of IL-1β and TNF-α for 3 days. (F, G) Differentiated astrocytes transfected with the indicated siRNAs were stimulated with IL-1β, TNF-α, and bFGF for 3 days. (D, F) Intracellular localization and semi-quantitative expression levels of Nrf2 were visualized by anti-Nrf2 immunostaining. Nuclei were visualized with DAPI. Scale bars: 50 μm. (E, G) The respective scatter dot plots show the relative intensities of nuclear Nrf2 determined by defining those in each control group as 1.0. The horizontal red lines indicate mean values (n > 500 cells per sample).

It has been well known that expression of *HO-1* is induced by Nrf2, a transcription factor responsible for stress responses (Alam *et al*, 1999; Loboda *et al*, 2016). We found that expression levels of *HO-1* in astrocytes stimulated with I/T/F were decreased by *Nrf2* knockdown (Fig. 3B and Appendix Fig. S3E). It has also been established that Nrf2 is constantly inactivated through proteasomal degradation in unstressed cells, and can be rapidly stabilized and activated in response to oxidative stress (Suzuki *et al*, 2019). Immunofluorescence analyses revealed that Nrf2 was expressed at low levels in cultured astrocytes under unstimulated conditions, but accumulated into the nuclei following stimulation with I/T, and the nuclear Nrf2 levels were further enhanced by stimulation with I/T/F (Fig. 3D and E). This nuclear accumulation of Nrf2 was strongly suppressed by *Ror2* knockdown and weakly suppressed by *Wnt5a* knockdown (Fig. 3F and G). Nrf2 can induce expression of a series of genes related to redox and iron homeostasis, in addition to *HO-1* (Tonelli *et al*, 2018). We found that *ferritin light chain 1* (*Flt1*), *glutamate-cysteine ligase modifier subunit* (*Gclm*), *solute carrier family 7 member 11* (*Slc7a11*), and *phosphogluconate dehydrogenase* (*Pgd*) were up-regulated significantly in reactive astrocytes induced by MCAO, but not LPS administration (Fig. EV3A). Moreover, expression levels of these genes were also up-regulated in *Ror2*- and *Nrf2*-dependent manners in cultured astrocytes stimulated with I/T/F (Fig. EV3B-D). These results indicate that increased expression of *Ror2* and subsequent activation of Wnt5a-Ror2 signaling might promote stabilization and nuclear accumulation of Nrf2, leading to transcriptional activation of its target genes.

### Wnt5a-Ror2 signaling prevents autophagic degradation of p62

We further investigated the molecular mechanism underlying Wnt5a-Ror2 signaling-mediated activation of Nrf2. Although Nrf2 has been known to be dissociated from its repressor Keap1 upon oxidative stress, thereby stabilized and accumulated in the nucleus (Suzuki *et al*., 2019), we found that cellular ROS levels were rather reduced in cultured astrocytes with strong nuclear accumulation of Nrf2 when stimulated with IL-1β, TNF-α, and bFGF (Fig. 4A). Alternative pathways for Nrf2 activation, that involve several Keap1 binding proteins, including p62 (also known as Sqstm1), are capable of inducing nuclear accumulation of Nrf2 through competitive interaction with Keap1 without its oxidative modification (Komatsu *et al*, 2010; Lau *et al*, 2010). Thus, we performed immunofluorescence staining of p62 in cultured astrocytes, and found that expression levels of p62 were increased by stimulation with I/T, and enhanced drastically by stimulation with I/T/F (Fig. 4B). Moreover, expression levels of p62 were decreased by knockdown of either *Ror2* or *Wnt5a*, but not *Nrf2* (Fig. 4C and D). In addition, intense nuclear accumulation of Nrf2 was observed in cells with higher expression levels of p62, and the nuclear Nrf2 intensity was decreased by suppressed expression of *p62* (Fig. 4E-G). It has been shown that phosphorylation of p62 at Ser-351 (Ser-349 in human) potentiates its binding capabilities with Keap1, leading to subsequent activation of Nrf2 (Ichimura *et al*., 2013). Consistently, puncta of phosphorylated p62 (P-p62) co-localized with Keap1 were observed clearly in cells stimulated with I/T/F, and were decreased by *Ror2* knockdown (Fig. 4H). Western blot analyses further confirmed that protein levels of p62 and P-p62, but not Keap1, were decreased when expression of *Ror2* was suppressed by siRNA (Fig. 4I). These results indicate that Wnt5a-Ror2 signaling increases the amounts of p62 phosphorylated at Ser-351, thereby promoting the nuclear accumulation of Nrf2 in astrocytes stimulated with I/T/F.

**Figure 4.**
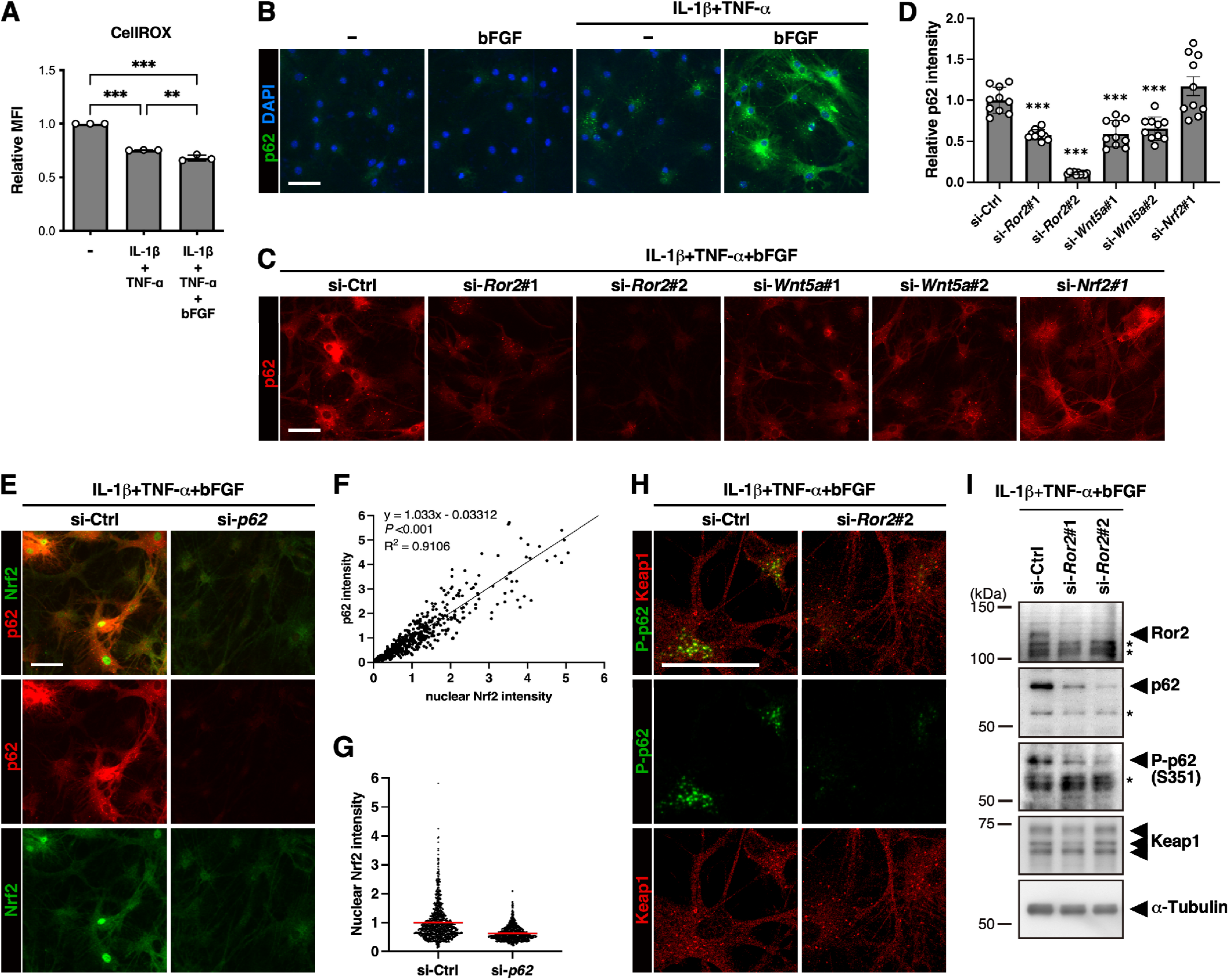
Wnt5a-Ror2 signaling enhances expression of p62. (A, B) Differentiated astrocytes were stimulated with IL-1β and TNF-α in the presence or absence of bFGF for 3 days. (A) The cellular ROS levels were determined by CellROX. The values are the mean ± SD of three independent samples. \*\**P*<0.01, \*\*\**P*<0.001 (Tukey’s test). (B) Expression levels of p62 were visualized by immunostaining. Nuclei were visualized with DAPI. Scale bar: 50 μm. (C-I) Differentiated astrocytes transfected with the indicated siRNAs were stimulated with IL-1β, TNF-α, and bFGF for 3 days. (C) Expression levels of p62 were visualized by immunostaining. Scale bar: 50 μm. (D) The bar graphs show the relative expression levels of p62 determined by defining those in si-Ctrl-transfected cells as 1.0. The values are the mean ± SD of 10 random fields. \**P*<0.05, \*\**P*<0.01, \*\*\**P*<0.001 (Dunnett’s test). (E) Expression levels of p62 (red) and Nrf2 (green) were visualized by immunostaining. Scale bar: 50 μm. (F) The scatter plot shows a positive correlation between expression levels of p62 and nuclear Nrf2. (G) The respective scatter dot plots show the relative intensities of nuclear Nrf2 determined by defining those in si-Ctrl-transfected cells as 1.0. The horizontal red lines indicate mean values (n > 700 cells per sample). (H) Expression levels of P-p62 (green) and Keap1 (red) were visualized by immunostaining. Scale bar: 50 μm. (I) Expression levels of the indicated proteins were analyzed by Western blotting. Asterisks indicate non-specific bands.

p62 has been known as a target gene for Nrf2 (Jain *et al*, 2010). However, protein levels of p62 were unaffected by suppressed expression of *Nrf2*, but decreased markedly by *Ror2* knockdown (Fig. 5A), suggesting that Ror2 might increase expression of p62 presumably through post-translational regulation. Cycloheximide chase analysis revealed that the half-life of p62 was indeed reduced by suppressed expression of *Ror2* (Fig. 5B and C). It has been shown that p62 acts as a selective autophagy receptor and is itself degraded by autophagy (Bjørkøy *et al*, 2005; Pankiv *et al*, 2007; Ichimura *et al*, 2008). Consistent with this finding, expression levels of p62 and LC3-II, another well-established substrate of autophagy, were increased when autophagy was inhibited by treatment with Bafilomycin A1 (BafA1) (Fig. 5D). In these autophagy-inhibited cells, nuclear Nrf2 levels were also markedly increased depending on increased expression of p62 (Fig. 5E-G). Interestingly, marked enhancement of p62 and nuclear Nrf2 levels induced by BafA1 was also observed in si-*Ror2*-treated cells at essentially identical levels as in si-Ctrl-treated cells (Fig. 5E-G). Unlike p62, LC3-II and neighbor of BRCA1 gene 1 (NBR1), another selective autophagy receptor that is structurally similar to p62, were unaffected by stimulation with I/T/F (Fig. EV4A). Consistently, expression levels of NBR1 were unaffected by suppressed expression of *Ror2*, while those were increased by treatment with BafA1 (Fig. EV4B and C)). In addition, BafA1 treatment markedly increased expression levels of p62 and nuclear Nrf2 even in unstimulated cells (Fig. EV4D-F), where expression of Ror2 remained at lower levels (Fig. EV4A). Collectively, these results indicate that p62 is continuously degraded by autophagy in cultured astrocytes, and that Wnt5a-Ror2 signaling, activated by stimulation with I/T/F, might selectively prevent autophagic degradation of p62 without affecting basal autophagy flux, thereby leading to activation of Nrf2 via intracellular accumulation of p62 (Fig. 5H).

**Figure 5.**
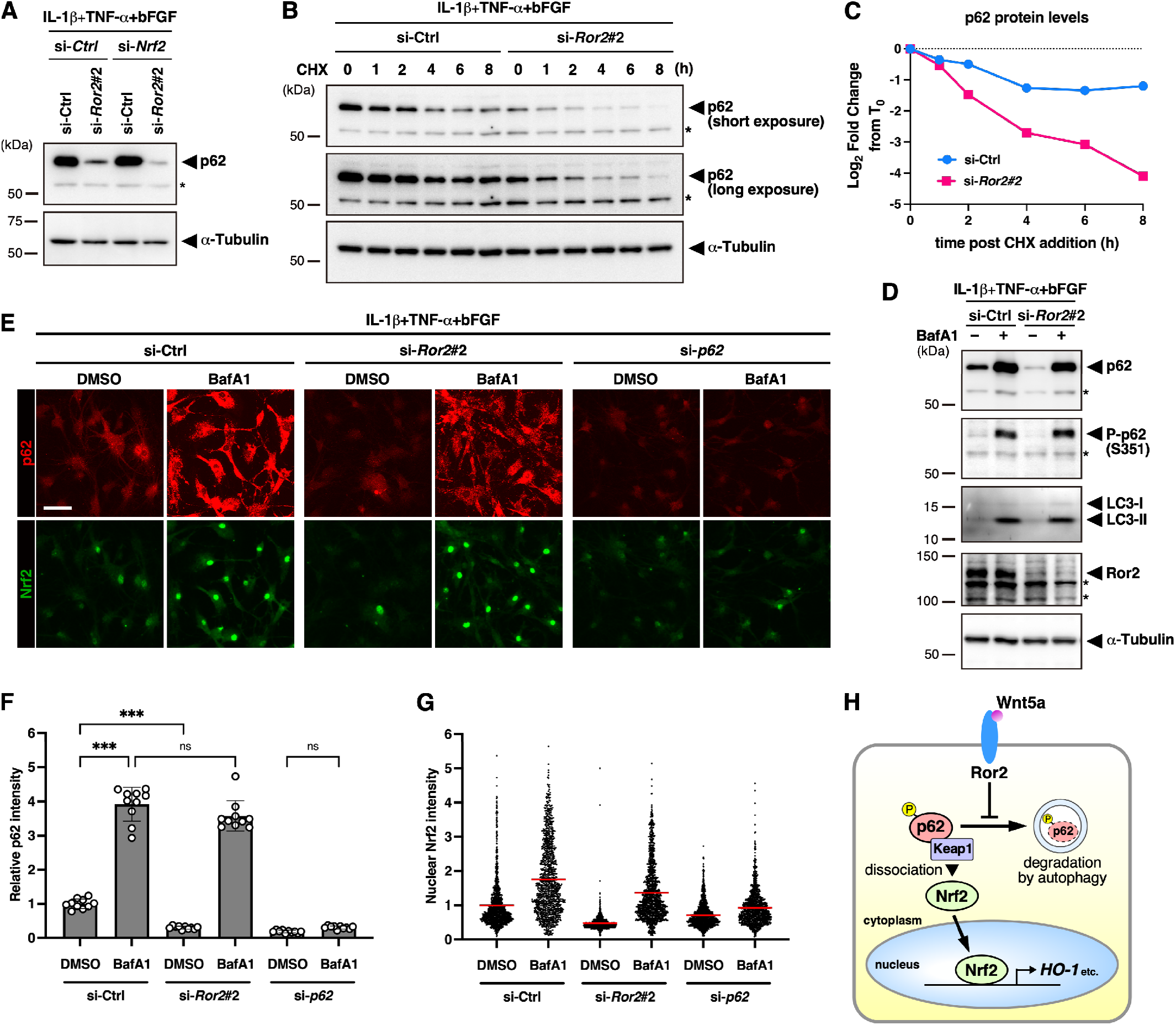
Wnt5a-Ror2 signaling prevents autophagic degradation of p62. (A) Differentiated astrocytes transfected with the indicated siRNAs were stimulated with IL-1β, TNF-α, and bFGF for 3 days. (B) Differentiated astrocytes transfected with *Ror2* siRNA (si-*Ror2*#2) or control siRNA (si-Ctrl) were stimulated with IL-1β, TNF-α, and bFGF for 3 days, and then treated with 20 μg/ml cycloheximide (CHX) for the indicated time (h). (A, B) Expression levels of p62 and α-Tubulin were analyzed by Western blotting. Asterisks indicate non-specific bands. (C) The relative band intensities of p62 were quantified and plotted on a semi-log graph. (D, E) Differentiated astrocytes transfected with the indicated siRNAs were stimulated with IL-1β, TNF-α, and bFGF for 2 days, then further treated with 100 nM Bafilomycin A1 (BafA1) or its vehicle (DMSO) alone for 24 h. (D) Expression levels of the indicated proteins were analyzed by Western blotting. Asterisks indicate non-specific bands. (E) Expression levels and intracellular distribution of p62 (red) and Nrf2 (green) were visualized by immunostaining. Scale bar: 50 μm. (F) The bar graphs show the relative expression levels of p62 determined by defining those in si-Ctrl-transfected and DMSO-treated cells as 1.0. The values are the mean ± SD of 10 random fields. \*\*\**P*<0.001 (Tukey’s test), ns: not significant. (G) The respective scatter dot plots show the relative intensities of nuclear Nrf2 determined by defining those in si-Ctrl-transfected and DMSO-treated cells as 1.0. The horizontal red lines indicate mean values (n > 1000 cells per sample). (H) A schematic representation of Nrf2 activation induced by Wnt5a-Ror2 signaling in astrocytes. Wnt5a-Ror2 signaling prevents autophagic degradation of p62, leading to intracellular accumulation of phosphorylated p62 that binds to Keap1 with high affinities. As a result, Nrf2, dissociated from Keap1, accumulates and translocates into the nucleus to induce expression of is target genes.

### Wnt5a-Ror2-p62-Nrf2 axis is activated in reactive astrocytes during tissue repair after intracerebral hemorrhage-induced brain injury

It has been reported that activation of Nrf2 pathway plays a critical role in promoting tissue protection following brain injuries with hemorrhage (Wang *et al*, 2007; Zhao *et al*, 2007). Thus, we next investigated whether Wnt5a-Ror2-p62-Nrf2 axis is involved in reactive astrocytes-mediated tissue protection and/or repair after ICH in early middle-aged mice. As observed in a SW injury model, increase in proliferating astrocytes and expression levels of *IL-1β*, *TNF-α*, *bFGF,* and *HB-EGF* was detected in the injured tissues in a collagenase-induced ICH mouse model (Appendix Fig. S4). Western blot analyses showed that expression levels of Ror2, but not Wnt5a, were up-regulated in the injured tissues on day 5 (D5) and 7 (D7), and returned to basal levels on day 14 (D14) after ICH (Fig. 6A). The electrophoretic mobility shift of Ror2 and increased phosphorylation levels of Dvl2, a well-known surrogate marker of canonical and noncanonical Wnt signaling, were observed on day 5 and 7 after ICH (Fig. 6A-C), indicating that Wnt5a-Ror2 signaling is indeed activated during these periods. Immunohistochemical analyses further demonstrated that Ror2 is expressed predominantly in GFAP-expressing reactive astrocytes surrounding the lesion core, where Iba1-positive inflammatory cells were densely accumulated (Fig. 6D). Any apparent differences in expression levels of p62 were not detected between ipsilateral and contralateral striatal brain hemispheres after the unilateral collagenase injection, while phosphorylation levels of p62 were significantly increased after day 5 of ICH (Fig. 6A-C). Interestingly, increased expression levels of P-p62 and Nrf2 were detected predominantly in Ror2-expressing reactive astrocytes rather than microglias on day 5 after ICH (Fig. 6E and F). Expression patterns and cell types of Pgd, an Nrf2 target gene product, were also similar to those of Ror2 (Fig. 6A-C and G). On the other hand, increased expression of HO-1 was induced from 1 day after ICH (Fig. 6A), and its strong expression was detected in Iba1-positive cells accumulating at the lesion core (Fig. 6H), presumably reflecting the fact that expression of *HO-1* can be induced independently of Nrf2 by hemin generated after erythrocyte lysis at the lesion following ICH as reported previously (Dunigan *et al*, 2018; Zhang *et al*, 2021).

**Figure 6.**
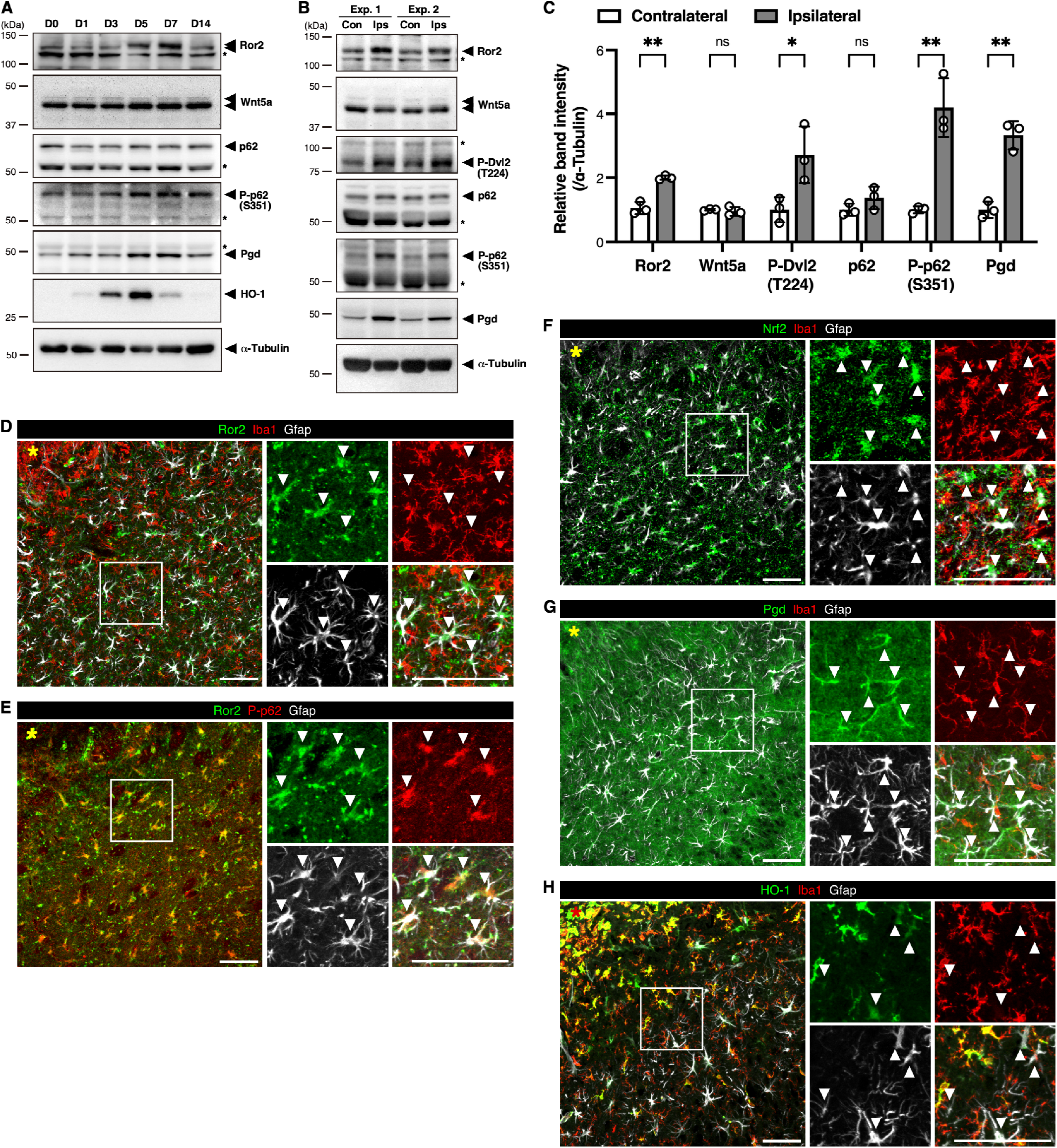
Wnt5a-Ror2-p62-Nrf2 axis is activated in reactive astrocytes following ICH in mice. (A, B) Adult male mice were subjected to ICH by intrastriatal injection of collagenase. (A) Expression levels of the indicated proteins in injured (D1, 3, 5, 7, 14) or uninjured (D0) striatal tissues were analyzed by Western blotting. Asterisks indicate non-specific bands. (B) Expression levels of the indicated proteins in ipsilateral (Ips) and contralateral (Con) striatal tissues on day 5 after ICH were analyzed by Western blotting. Data from two independent samples are shown. Asterisks indicate non-specific bands. (C) The bar graphs show the relative band intensities of the indicated proteins normalized to those of α-Tubulin. Data are shown as the mean ± SD (n = 3 animals per group). \**P*<0.05, \*\**P*<0.01. (multiple Student’s *t*-test), ns: not significant. (D-H) Striatal tissues on day 5 after ICH were immunostained with the indicated antibodies. The right panels are magnified views of the square in the left panel. Arrowheads indicate GFAP-expressing reactive astrocytes. Asterisks indicate the injury sites. Scale bars: 100 μm.

Hemin is highly cytotoxic and can cause secondary brain damage following ICH (Robinson *et al*, 2009). We found that cultured astrocytes stimulated with I/T/F were more resistant to hemin-mediated cytotoxicity than untreated cells (Fig. 7A). Resistance of these astrocytes to hemin was suppressed by *Ror2* knockdown (Fig. 7B) and enhanced by treatment with recombinant Wnt5a (Fig. 7C). Analyses of intracellular Fe^2+^ and lipid peroxidation with the fluorescence probes FerroOrange and Liperfluo revealed that both levels were increased within 2 h after hemin treatment and further increased when *Ror2* expression was suppressed (Fig. 7D-G). These results suggest that activation of Wnt5a-Ror2 signaling prevents cell death elicited by an excess iron-dependent accumulation of lipid peroxides in hemin-exposed astrocytes.

**Figure 7.**
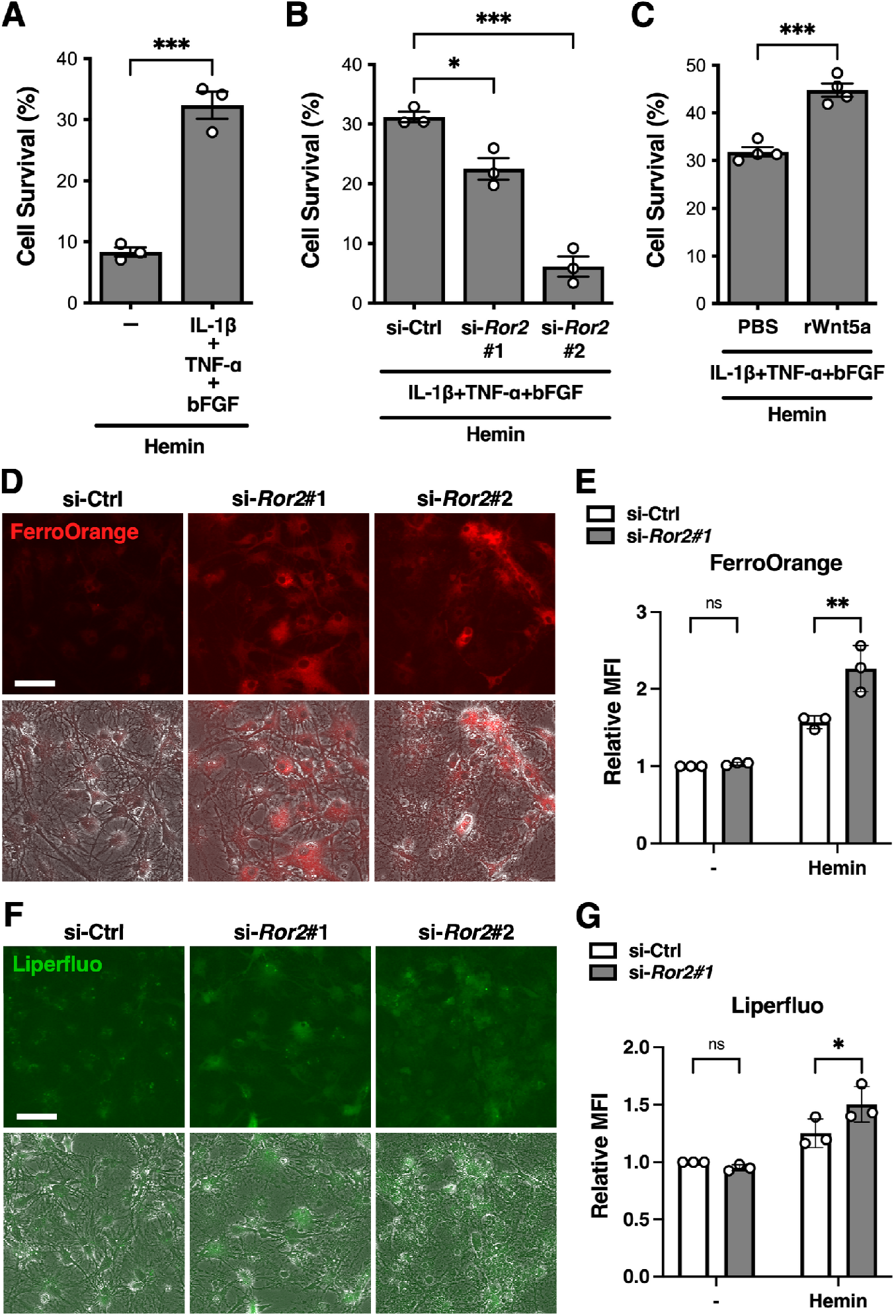
Wnt5a-Ror2 signaling protects astrocytes from hemin-induced cytotoxicity. (A-C) Differentiated astrocytes were treated with or without 10 μM hemin for 6 h, and subjected to cell survival analysis. The respective bar graphs show percentages of cell survival following hemin treatment determined by defining the viability of hemin-untreated cells in each group as 100 %. The values are the mean ± SD of three independent experiments. \**P*<0.05, \*\*\**P*<0.001 (Student’s *t*-test for A, C; Dunnett’s test for B). Astrocytes were cultured as below prior to hemin treatment. (A) Cells were cultured with or without IL-1β, TNF-α, and bFGF for 3 days. (B) Cells transfected with the indicated siRNAs were cultured with IL-1β, TNF-α, and bFGF for 3 days. (C) Cells were cultured with IL-1β, TNF-α, and bFGF together with recombinant Wnt5a (rWnt5a) or its vehicle (PBS) for 3 days. (D, E) Differentiated astrocytes transfected with the indicated siRNAs were cultured with IL-1β, TNF-α, and bFGF for 3 days, followed by treatment with or without 10 μM hemin for 2 h. Levels of intracellular ferrous iron (Fe^2+^) (D) and lipid peroxide (E) were visualized by FerroOrange (red) and Liperfluo (green), respectively. Most of the cells transfected with si-*Ror2*#2 were already dead at 2 h after hemin treatment, but the surviving cells showed intense signals of FerroOrange and Liperfluo. Scale bars: 50 μm. (E, G) The respective bar graphs show the relative mean fluorescent intensities (MFIs) of FerroOrange (E) or Liperfluo (G) staining in the indicated cells determined by defining the MFI in hemin-untreated si-Ctrl-transfected cells as 1.0. The values are the mean ± SD of three independent experiments. \**P*<0.05, \*\**P*<0.01 (Tukey’s test), ns: not significant.

It has also been reported that excessive amounts of hemin and Fe^2+^ induce dysfunction of the BBB (Imai *et al*, 2019). In fact, on day 5 after ICH, leakage of exogenous FITC-dextran tracers and endogenous plasma immunoglobulin G (IgG) was detected in the vicinity of reactive astrocytes (Fig. EV5A and B. In addition, amounts of leaked IgG heavy and light chains (IgG-H and IgG-L) and expression levels of HO-1 gradually decreased around day 7 of ICH (Fig. 6A and Fig. EV5C), suggesting that activation of Wnt5a-Ror2 signaling in reactive astrocytes might restore the integrity of the BBB along with heme clearance. To clarify whether Wnt5a-Ror2 signaling indeed activates p62-Nrf2 axis in reactive astrocytes and regulates the BBB integrity, we prepared an AAV vector capable of transducing Wnt5a exogenously in an astrocyte-specific manner.

Overexpression of Wnt5a induced the electrophoretic mobility shift of Ror2 via its hyperphosphorylation and increased expression levels of p62 in cultured astrocytes stimulated with I/T/F, but not in untreated cells (Fig. 8A). Furthermore, the mobility shift of Ror2 and modestly increased P-p62 and Pgd levels were observed in the injured tissues on day 5 after ICH in AAV-Wnt5a-injected mice compared with AAV-GFP-injected ones (Fig. 8B and C), although expression levels of Gfap and Aldh1l1, astrocyte markers, were unaffected (Fig. 8B and C). In contrast, expression levels of HO-1 and amounts of IgG-H, L in the injured tissues were decreased by overexpression of Wnt5a (Fig. 8B and C). These results indicate that Wnt5a-Ror2 signaling can activate p62-Nrf2 axis in reactive astrocytes and promote BBB repair possibly via heme and iron clearance.

**Figure 8.**
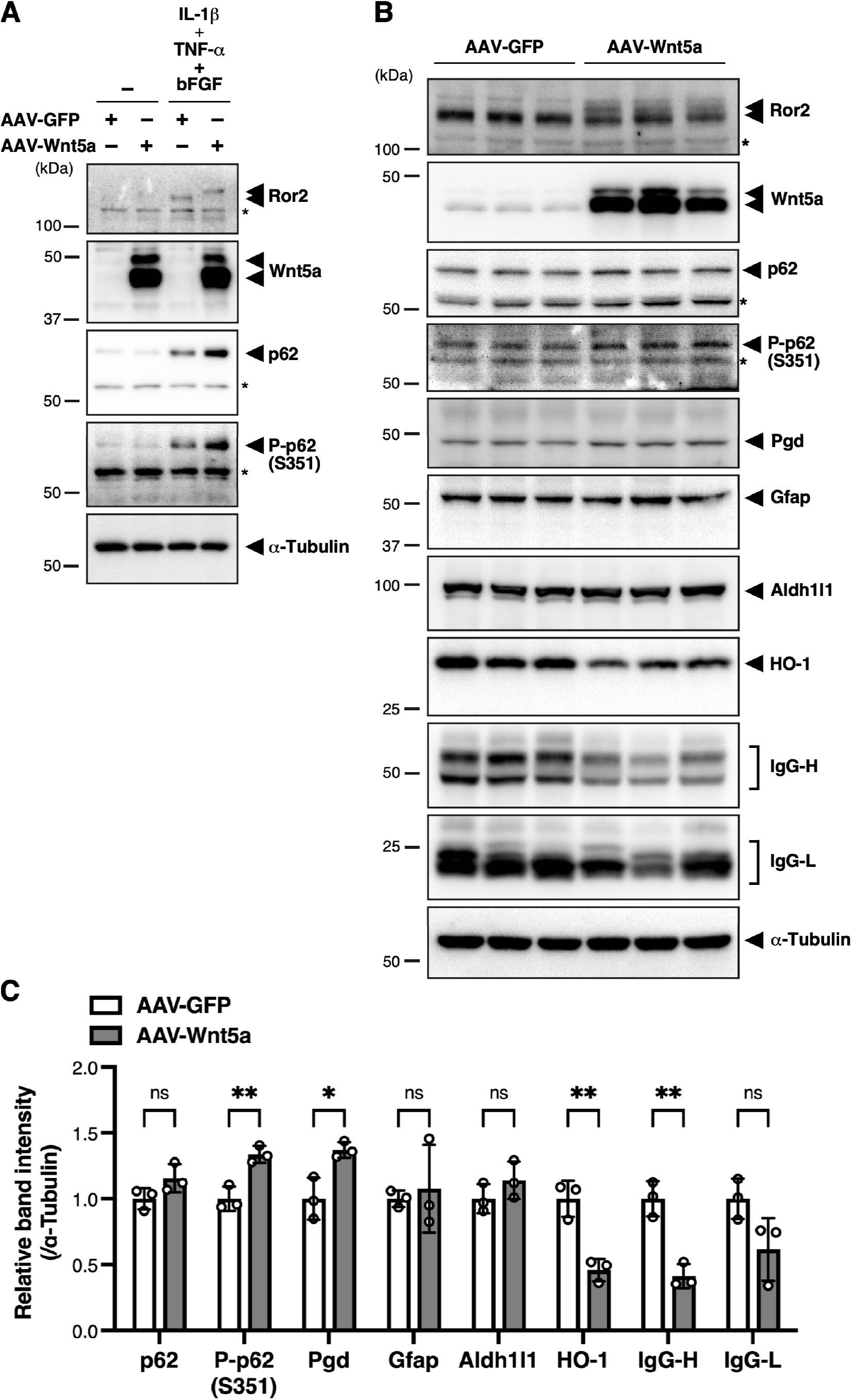
Overexpression of Wnt5a in astrocytes results in increased amount of phosphorylated p62 and reduced extravasation of IgG in mouse brains after ICH. (A) Differentiated astrocytes infected with either AAV-GFP or AAV-Wnt5a were cultured in the presence or absence of IL-1β, TNF-α, and bFGF for 3 days. Expression levels of the indicated proteins were analyzed by Western blotting. Asterisks indicate non-specific bands. (B) Adult male mice, previously injected with astrocyte-specific AAV-GFP or AAV-Wnt5a in the striatum, were subjected to ICH by intrastriatal injection of collagenase. Expression levels of the indicated proteins in the injured striatal tissues on day 5 after ICH were analyzed by Western blotting. Data from three independent samples are shown for each group. Asterisks indicate non-specific bands. (C) The bar graphs show the relative band intensities of the indicated proteins normalized to those of α-Tubulin. Data are shown as the mean ± SD (n = 3 animals per group). \**P*<0.05, \*\**P*<0.01. (multiple Student’s *t*-test), ns: not significant.

## Discussion

Reactive astrocytes exhibit diverse functions in a context-dependent manner, that are based primarily on gene expression patterns induced by different types of transcription factors. Under pathological conditions, including brain injury, reactive astrocytes are often observed in close proximity to activated microglias and/or infiltrated macrophages. Thus, proinflammatory cytokines produced by activated microglias/macrophages may be involved in regulating the functions of transcription factors that regulate context-dependent functions of reactive astrocytes. In this study, we focused on the fact that bFGF and HB-EGF, which act as mitogens for astrocytes, are produced in the injured brains, and found that stimulation of astrocytes by these growth factors together with inflammatory cytokines results in synergistic enhancement of E2F1-mediated transcriptional induction of *Ror2*. We have previously shown that Ror2 can promote E2F1-mediated cell cycle progression of fibroblasts stimulated with bFGF. On the other hand, it has been reported that E2F1-mediated cell cycle progression can be inhibited by co-stimulation with inflammatory cytokines in an NF-κB-dependent manner (Araki *et al*, 2008). Consistent with this report, I/T indeed inhibited cell cycle progression of cultured astrocytes stimulated with bFGF although E2F1-mediated expression of *Ror2* was enhanced in these cells, suggesting that Ror2 might exhibit functions other than regulating cell cycle progression in astrocytes stimulated with growth factors together with inflammatory cytokines. We also performed gene expression analyses, including RNA-Seq, using cultured astrocytes stimulated with I/T/F to explore novel functions of Ror2 in reactive astrocytes in the injured brains, and found that Wnt5a-Ror2 signaling is involved in p62-dependent activation of Nrf2, conferring protection against oxidative damage in astrocytes. These findings indicate that growth factors including bFGF produced in the injured brains promote not only the proliferation of astrocytes located near the injured area, but also the function of reactive astrocytes contributing to tissue protection and repair following injury. Our RNA-Seq analysis also provides evidence that growth factor signaling and inflammatory cytokine signaling act cooperatively to up-regulate expression of multiple genes in addition to Nrf2 target genes in a Ror2-dependent or - independent manner. Future study will be required to clarify the roles of these genes in reactive astrocytes in regulating tissue repair following brain injury.

It is well established that p62 acts as a receptor for selective autophagy and promotes the degradation of misfolded, ubiquitinated proteins (Wurzer *et al*, 2015; Zaffagnini *et al*, 2018). After binding with ubiquitinated proteins, p62 undergoes phase separation to form liquid droplets, which are subsequently wrapped into autophagosomes followed by their degradation (Komatsu, 2022). It has also been shown that p62-droplets serve as a signaling hub for Nrf2 activation through the interaction with Keap1 in a phosphorylation-dependent manner, and that suppression of autophagy leads to accumulation of phosphorylated p62 and persistent activation of Nrf2 (Ichimura *et al*., 2013). Assuming a negative correlation between p62-mediated autophagic degradation and activation of Nrf2, it is conceivable that the molecular mechanisms determining either function to be activated depend on cellular contexts. In this respect, our findings show for the first time that Wnt5a-Ror2 signaling promotes activation of p62-Nrf2 pathway by preventing autophagic degradation of p62 without affecting the basal autophagic activities. How does Wnt5a-Ror2 signaling selectively prevent autophagic degradation of p62? NBR1 has been shown to interact with p62 and prevent its autophagic degradation, leading to stabilization of p62 and activation of Nrf2 (Sánchez-Martín *et al*, 2020). However, expression levels of NBR1 were unaffected by *Ror2* knockdown, indicating that Wnt5a-Ror2 signaling regulates p62 stability independently of NBR1. p62-droplets interact with Autophagy-related gene (Atg) proteins to form isolation membranes around them, leading to their subsequent degradation by autophagy (Kageyama *et al*, 2021), although the molecular mechanisms determining the initiation and progression of this process remain unknown. It can be assumed that Wnt5a-Ror2 signaling might regulate p62 stability by inhibiting or delaying the interaction of p62 with Atg proteins. Further study is required to clarify how Wnt5a-Ror2 signaling promotes the stability of p62.

Autophagy plays an important role in promoting cell survival under hypoxic conditions in the injured brains with vascular damage. On the other hand, hemin released from hemoglobin during hemorrhage at these injury sites causes cellular damage due to oxidative stress. Therefore, Wnt5a-Ror2 signaling may play an important role in protecting cells from hemin-induced oxidative stress in the injured brains by robustly activating Nrf2 under autophagy-activating conditions. Our present findings provide evidence, supporting a notion that Wnt5a-Ror2-p62-Nrf2 signaling axis is activated predominantly in reactive astrocytes surrounding the injured areas, but not in activated microglias, indicating that this signaling axis might play a context-dependent and cell type-specific function. It has been reported that activation of Nrf2 in astrocytes can protect themselves and neighboring cells, including neurons, from oxidative damages in the injured or diseased brains and spinal cords, thereby improve pathological conditions (Vargas *et al*, 2008; Draheim *et al*, 2016; Sigfridsson *et al*, 2018; Zhao *et al*, 2022). Consistent with these reports, we found that activation of p62-Nrf2 pathway by Wnt5a-Ror2 signaling activated in astrocytes reduces hemin-induced cellular damage *in vitro* and promotes repair of the disrupted BBB *in vivo*. It will be of interest to examine the possibility that Wnt5a-Ror2-p62-Nrf2 signaling axis might also play an important role in promoting tissue repair following injury in other tissues. It can be envisaged that Wnt5a-Ror2-p62-Nrf2 signaling axis might be a suitable target for the development of novel therapeutic approaches to promote tissue repair properly after injury with vascular disruption, especially in the brains.

## Materials and Methods

### AAV vectors and virus production

We constructed *pAAV-hGFAP-Lck-eGFP* and *pAAV-hGFAP-Lck-eGFP-T2A-Wnt5a* as AAV transfer plasmids through PCR-based subcloning into *pAAV-hGFAP* vector, in which the *CMV* promoter of *pAAV-CMV* vector (Takara Bio) was replaced by *gfaABC1D*, an astrocyte-specific minimal human *GFAP* promoter (Lee *et al*, 2008). *Lck-eGFP* indicates the gene encoding a membrane-targeted form of eGFP. AAV-PHP.eB viruses were packaged in AAVpro 293T cells (Clontech). Briefly, AAVpro 293T cells were cultured in DMEM (FUJIFILM Wako Pure Chemical) supplemented with 10% (v/v) FBS (NICHIREI) and transfected with the *pUCmini-iCAP-PHP.eB* plasmid (Addgene), *pHelper* plasmid (Takara Bio) and AAV transfer plasmids using TransIT-VirusGEN Transfection reagent (Mirus). The next day, transfection medium was replaced with fresh medium, and further cultured for 2 days. The culture medium was collected 72 h after transfection and concentrated using AAVanced Concentration Reagent (System Biosciences) according to the manufacturer’s protocol. Viral titer was measured using AAVpro Titration Kit (for Real Time PCR) Ver. 2 (Takara Bio).

### Animals and Stereotaxic AAV injection

In this study, we used young (2–3 months-old) and early middle-aged (8–10 months-old) male C57BL/6N mice (Japan SLC, Inc.) for the stab-wound injury experiments and ICH model experiments, respectively. All animal experiments were approved by the Institutional Animal Care and Use Committee (Permission numbers: P191017 and P210403-R1) and carried out according to the Kobe University Animal Experimentation Regulations. For the AAV-mediated gene transfer experiments, mice were anesthetized with isoflurane (FUJIFILM Wako Pure Chemical), and then 0.25 μL of AAV solution (1 x 10^13^ vg/mL) was injected twice into the right striatum (0.5 mm anterior to bregma, 2 mm lateral from the midline) at two depths (3 mm and 4 mm from the skull) at a flow rate of 0.1 μL/min using a Hamilton syringe driven by a syringe pump. Following the injection, the needle was left in place for 10 min before removal. Two weeks after AAV injection, mice were used for the ICH model experiments.

### Surgical procedures

Stab-wound injury experiments were performed as previously described (Endo *et al*., 2017). Briefly, mice were anesthetized with isoflurane and subjected to a stab wound injury (1 mm in depth and 1.5–2 mm in length) on the right cortical hemisphere using an ophthalmic knife and stereotactic apparatus. For the ICH model experiments, mice were anesthetized with isoflurane, and then collagenase VII-S (0.03U in 0.5 μL saline, Sigma-Aldrich) was injected into the right striatum (0.5 mm anterior to bregma, 2 mm lateral from the midline, and 3.5 mm in depth) at a flow rate of 0.1 μL/min using a Hamilton syringe driven by a syringe pump. Following the injection, the needle was left in place for 10 min before removal.

### Fluorescence-activated cell sorting

Mice were anesthetized with isoflurane and transcardially perfused with ice-cold PBS. The brains were immediately dissected and the injured and uninjured neocortices were punched out (Ø 0.2 cm). Tissues were then enzymatically digested with papain solution [8 units/mL papain (Worthington), 20 μg/mL DNase I (Sigma-Aldrich), 0.2 mg/mL L-cysteine hydrochloride (Sigma-Aldrich), 0.5 mM EDTA in EBSS (Life Technologies) equilibrated with sterile 95% O_2_ and 5% CO_2_] for 30 min at 34℃, and mechanically dissociated in FACS buffer [FluoroBriteDMEM (Thermo Fisher Scientific) containing 2 mM EDTA and 0.5% (w/v) BSA] with DNase I to obtain single-cell suspensions. Each cell suspension was then passed through a 40 μm cell strainer, and subjected to debris removal and red blood cell removal using debris removal solution (Miltenyi Biotec) and red blood cell lysis solution (Miltenyi Biotec), respectively, according to the manufacturer’s instructions. Cells were resuspended in FACS buffer containing FcR blocking reagent (Miltenyi Biotec), and incubated for 10 min at 4℃, followed by incubation with BV421-conjugated anti-CD45 (BD Biosciences, 563890, 1:100), PE-conjugated anti-O4 (Miltenyi Biotec, 130-117-507, 1:50), and APC-conjugated anti-ACSA-2 (Miltenyi Biotec, 130-116-245, 1:100) antibodies for 30 min at 4℃. Unlabeled cells were used as a negative control. Prior to FACS sorting, the cells were washed twice and resuspended in FACS buffer containing 7-AAD viability staining solution (Life Technologies). Cell sorting was performed using a BD FACSAria III (BD Biosciences) equipped with a 100 μm nozzle. Cells were sorted directly into TRIzol LS reagent (Thermo Fisher Scientific).

### Immunohistochemistry

Mice were anesthetized with isoflurane and transcardially perfused with 4% (w/v) paraformaldehyde (PFA). For the BBB permeability analysis, 10 mg/mL FITC-dextran solution (10,000 Da, Tokyo Chemical Industry) was perfused prior to perfusion with PFA as described previously (Natarajan *et al*, 2017). The brains were immediately dissected and post-fixed with 4% PFA for 2 h at room temperature (RT) or overnight at 4℃. After equilibration with 30% (w/v) sucrose in PBS, the fixed brains were embedded in OCT compound (Sakura Finetek) and frozen. Coronal sections were prepared by cutting the frozen brains with a cryostat at a thickness of 20 μm. Brain sections were treated with 1% (v/v) hydrogen peroxide in PBS for 1 h to quench endogenous peroxidase activity and permeabilized with 0.2% (v/v) Triton X-100 in PBS for 20 min. After blocking with the normal 2.5% (v/v) serum matching the species of the secondary antibody, the sections were incubated with the respective antibodies as follows: mouse anti-GFAP (Thermo Fisher Scientific, 14-9892-82, 1:1000), rabbit anti-S100β (Agilent, 0311, ready to use), rabbit anti-Iba1 (FUJIFILM Wako Pure Chemical, 019-19741, 1:500), rat anti-Ki67 (Thermo Fisher Scientific, 14-5698-82, 1:300), goat anti-Ror2 (R&D Systems, AF2064, 1:100), rabbit anti-HO-1 (Abcam, ab13243, 1:1000), rabbit anti-phospho-p62 (Ser-351) (MBL, PM074, 1:5000), rabbit anti-Nrf2 (Cell Signaling Technology, 12721, 1:500), and rabbit anti-Pgd (Proteintech, 14718-1-AP, 1:500). For staining with mouse monoclonal anti-GFAP antibody, M.O.M. Blocking Reagent (Vector Laboratories) was used to reduce endogenous mouse IgG staining. The following secondary antibodies were used: Alexa Fluor 546 goat anti-rabbit IgG (Thermo Fisher Scientific, A11035, 1:500), Alexa Fluor 635 goat anti-mouse IgG (Thermo Fisher Scientific, A31575, 1:500), and CF488A donkey anti-rat IgG (Biotium, 20027, 1:500). For multiple immunofluorescence staining, the Tyramide Signal Amplification (TSA) method was used to detect Ror2, phospho-p62, Nrf2, and Pgd. At the secondary antibody labeling step, sections were incubated with the appropriate ImmPRESS HRP polymer-conjugated secondary antibodies (anti-goat or anti-rabbit IgG, Vector Laboratories) for 1 h and then stained using the TSA Kit with Alexa Fluor 488 tyramide reagent (Life Technologies), according to the manufacturer’s instructions.

### Cell culture, transfection, and infection

Neural progenitor cells (NPCs)-derived astrocytes were prepared as described previously (Endo *et al*., 2017). In brief, NPCs were isolated from cerebral neocortices of neonatal (P0) ICR mice and cultured as neurospheres in NPC medium [DMEM/F12 (FujiFilm Wako Pure Chemical) supplemented with Neurobrew-21 (Miltenyi Biotec), 20 ng/mL bFGF (FujiFilm Wako Pure Chemical), and 20 ng/mL EGF (FujiFilm Wako Pure Chemical)]. The primary neurospheres were dissociated into a single-cell suspension using Accutase (Nacalai Tesque) and seeded in poly-L-lysine (Sigma-Aldrich)-and Matrigel Matrix (Corning)-coated dishes and subsequently passaged every 3 days with fresh NPC medium. To induce differentiation into astrocytes, NPCs were cultured on poly-L-lysine-coated dishes in differentiation medium [DMEM/F12 supplemented with Neurobrew-21 and 2 % (v/v) FBS]. After 3 days in culture, cells were treated with 10 μM cytosine β-D-arabinofuranoside (Sigma-Aldrich) for 5 days to deplete oligodendrocyte progenitor cells. After 10-12 days in culture, the cells were used as differentiated astrocytes. Following differentiation, culture medium was replaced with serum-free DMEM/F12 supplemented with Neurobrew-21, and then stimulated with 20 ng/mL bFGF or 20 ng/mL HB-EGF (PeproTech) in the absence or presence of 10 ng/mL IL-1β (FujiFilm Wako Pure Chemical) and 20 ng/mL TNF-α (PeproTech). Differentiated astrocytes expressing ERT2-E2F1-1′C were prepared from NPCs that were previously infected with retrovirus encoding *ERT2-E2F1-1′C* followed by selection with 0.25 μg/mL puromycin (Nacalai tesque). For the siRNA-transfection experiments, differentiated astrocytes were transfected with the respective siRNA oligonucleotides by Lipofectamine RNAiMax reagent (Thermo Fisher Scientific). Silencer select siRNAs targeting moue *Ep400* (si-*Ep400*#1, s93640; si-*Ep400*#2, s93641), *Ror2* (si-*Ror2*#1, s77263; si-*Ror2*#2, s77265), *Ror1* (si-*Ror1*#1, s77260; si-*Ror1*#2, s77261), *Wnt5a* (si-*Wnt5a*#1, s76087; si-*Wnt5a*#2, s76088), *Nrf2* (si-*Nrf2*#1, s70521; si-*Nrf2*#2, s70523), and *p62* (si-*p62*, s71144) and their control siRNA (Silencer select negative control No. 1) were purchased from Thermo Fisher Scientific. The sequences of the siRNAs used are listed in Appendix Table S1. For the AAV-mediated gene transfer experiments, NPCs were cultured overnight on poly-L-lysine-coated dishes in differentiation medium, and then treated with AAV solution (1 x 10^10^ vg/mL) for 2 days and further differentiated as above. Two weeks after AAV infection, differentiated astrocytes were used for the experiments.

### Cell survival analysis

Cells were washed once with PBS and cultured in non-supplemented DMEM/F12 with or without 10 μM hemin (FujiFilm Wako Pure Chemical) for 6 h. Cell survival rate was determined using Cell Counting Kit-8 (Dojindo)

### Quantitative RT-PCR

Total RNAs were isolated from tissues or cells using Sepasol-RNA I Super G (Nacalai Tesque) or TRIzol LS reagent. cDNAs were synthesized from these RNAs as templates using PrimeScript RT Reagent (Takara Bio). Expression levels of the respective genes of interest were measured using LightCycler 480 II system (Roche). The amounts of the respective mRNAs were normalized relative to those of *18S ribosomal RNA*. The primers used are listed in Appendix Table S2.

### Chromatin Immunoprecipitation (ChIP) Assay

ChIP assay was performed using SimpleChIP Enzymatic Chromatin IP Kit (Cell Signaling Technology) according to the manufacturer’s instructions. Briefly, cells were fixed with 1 % (w/v) PFA for 10 min at RT, followed by incubation with glycine. After washing with ice-cold PBS, cells were digested with micrococcal nuclease. The nuclei were sonicated for 4 cycles (sonication, 20 seconds; on ice, 60 seconds) by a cell disruptor (UD-201, Tomy) and subjected to immunoprecipitation with anti-E2F1 antibody (Santa Cruz Biotechnologies, sc-251) or control mouse IgG (Jackson ImmunoResearch Laboratories, 015-000-003). After decrosslinking and purification of the DNA fragments, the amounts of the DNA fragments of interest were measured by LightCycler 480 II System with a primer pair (5’-CAAGGGACGCCTAGTTAATG-3’ and 5’-GACTTCTAGGCACAAAGGTG-3’) for *Ror2* promoter and normalized relative to those in input materials.

### Western blotting

To prepare whole cell lysates, cells were solubilized with and tissues were homogenized in lysis buffer containing 50 mM Hepes (pH 7.5), 150 mM NaCl, 1% (v/v) Nonidet P-40, 1 mM EDTA, 10 μg/ml leupeptin, and 10 μg/ml aprotinin, 1 mM p-APMSF, and PhosSTOP phosphatase inhibitor cocktail (Sigma-Aldrich) and removed insoluble materials by centrifugation. Proteins were separated by SDS-PAGE (7.5%–10% PAG) and transferred onto Immobilon-P membranes (Merck Millipore). Membranes were immunoblotted with the respective antibodies as follows: rabbit anti-Ror2 (Kani *et al*, 2004), rabbit anti-Wnt5a (Cell Signaling Technology, 2530, 1:1000), mouse anti-α-Tubulin (Sigma-Aldrich, CP06, 1:2000), rabbit anti-p62 (MBL, PM045, 1:2000), rabbit anti-phospho-p62 (S-351) (MBL, PM074, 1:1000), mouse anti-LC3 (MBL, PM036, 1:1000), rabbit anti-Keap1 (Proteintech, 10503-2-AP, 1:1000), rabbit anti-NBR1 (Cell Signaling Technology, 20145, 1:1000), mouse anti-Pgd (Santa Cruz Biotechnology, 398977, 1:100), rabbit anti-HO-1 (Abcam, ab13243, 1:1000), rabbit anti-phospho-Dvl2 (T224) (Abcam, ab124941, 1:1000), mouse anti-GFAP (Thermo Fisher Scientific, 14-9892-82, 1:1000), and rabbit anti-Aldh1l1 (Abcam, ab87117, 1:1000). The following secondary antibodies were used: goat anti-rabbit IgG HRP conjugate (Bio-Rad, 170-6515, 1:5000) and goat anti-mouse IgG HRP conjugate (Bio-Rad, 170-6516, 1:5000). Immunoreactive bands were visualized by using Western Lightning Plus-ECL (Perkin Elmer) or ImmunoStar LD (Fujifilm Wako Pure Chemical), and detected using the ECL detection system (LAS-1000; FujiFilm Wako Pure Chemical). The relative band intensities were measured using ImageJ software (National Institutes of Health).

### Immunofluorescence analysis

Cells were fixed with 4 % (w/v) PFA for 20 min at RT, followed by treatment with 1% (v/v) hydrogen peroxide in PBS for 1 h. After treating with blocking buffer [3% (w/v) BSA and 0.2% (v/v) Triton X-100 in PBS], the cells were incubated with the respective antibodies as follows: rabbit anti-Nrf2 (Cell Signaling Technology, 12721, 1:1000), rabbit anti-p62 (MBL, PM045, 1:2000), rabbit anti-phospho-p62 (Ser-351) (MBL, PM074, 1:5000), and rabbit anti-Keap1 (Proteintech, 10503-2-AP, 1:200). Alexa Fluor 488 Tyramide SuperBoost Kit, goat anti-rabbit IgG (Thermo Fisher Scientific, B40922) was used to detect Nrf2 and phospho-p62, which allows multiple labeling with primary antibodies raised in the same species. Alexa Fluor 546 goat anti-rabbit IgG (Thermo Fisher Scientific, A11035, 1:500) was used to detect p62 and Keap1, and the nuclei were stained with DAPI.

### RNA-Seq and bioinformatics analysis

Total RNAs were extracted from cells using the RNeasy Mini kit (Qiagen). AmpliSeq libraries were constructed using the Ion AmpliSeq Transcriptome Mouse Gene Expression Kit (Thermo Fisher Scientific) that is designed for targeted amplification of over 20,000 mouse RefSeq genes simultaneously in a single primer pool, and then barcoded using Ion Xpress Barcode Adapters (Thermo Fisher Scientific). For each sample, 100 ng of total RNA was used for cDNA library preparation. Barcoded libraries were clonally amplified using the Ion OneTouch 2 System (Thermo Fisher Scientific) and then sequenced on an Ion Torrent PGM (Thermo Fisher Scientific). RNA-seq reads were mapped to the reference genome (mouse: GRCm38) and differential expression analysis was performed using CLC bio Genomics Workbench Version 12.0 (CLC bio). Genes with *q*-value (FDR-adjusted *P*-value) ≤ 0.05 and fold change < 1 or > 1 were defined as down-regulated or up-regulated differentially expressed genes (DEGs), respectively. We first extracted DEGs that were up-regulated by IL-1β stimulation and further enhanced by co-stimulation with bFGF. Among them, we further extracted DEGs that were down-regulated by *Ror2* knockdown in cells stimulated with IL-1β and bFGF. Among the genes identified by the RNA-Seq analysis, we finally extracted genes commonly up-regulated in reactive astrocytes in the injured brains following MCAO and SW by analyzing microarray gene expression data from Gene Expression Omnibus GSE35338 and GSE66370 (Table EV1).

### Data Presentation and Statistics

Fluorescence images of tissues or cultured cell were obtained using a laser scanning confocal imaging system (LSM710; Carl Zeiss) or a fluorescence microscope (BZ-X700; Keyence), respectively, and processed using ImageJ and GIMP 2.10 software (https://gimp.org). The relative fluorescence intensities were measured using ImageJ or hybrid cell count software (Keyence). Statistical analyses were performed using the GraphPad Prism 9.0 (GraphPad Software). Significance was determined as \**P* < 0.05, \*\**P* < 0.01, or \*\*\**P* < 0.001 compared with control using Student’s *t*-test when two groups were compared or using one-way ANOVA followed by Dunnett’s or Tukey’s post hoc test when more than three groups were analyzed.

### Data availability

RNA-Seq raw data files were deposited at the DDBJ Sequenced Read Archive under the accession number DRA016030.

## Acknowledgements

This work was supported by grants from Japan Agency for Medical Research and Development (AMED) (JP23gm1210005) to YM, and MEXT/JSPS KAKENHI (JP18K06486) to ME, and by COI-NEXT support Unit for Imaging Science at Kento (JPMJPF2018) to YM. This work also received funding to ME from the collaborative research project program supported by Sumitomo Pharma Co., Ltd.

## Author contributions

ME and YM designed research; ME, YT, MF, and HS performed experiments; ME analyzed data; ME and YM wrote the manuscript.

## Expanded View Figures

**Figure EV1. Expression of *E2F1* is up-regulated through the synergistic action of bFGF and inflammatory cytokines in cultured astrocytes irrespective of their cell cycle progression.** Differentiated astrocytes were stimulated with either IL-1β and TNF-α or vehicle alone in the presence or absence of bFGF for 3 days. Expression levels of *E2F1*, *CyclinE1*, and *Ki67* were analyzed by qRT-PCR. Relative expression values were determined by defining the expression level of the indicated genes in untreated cells (-) as 1.0. The values are the mean ± SD of three independent experiments. \**P*<0.05, \*\**P*<0.01, \*\*\**P*<0.001 (Tukey’s test), ns: not significant.

**Figure EV2. HO-1 is expressed highly in Ror2-expressing proliferative reactive astrocytes in the injured brains.** (A) Neocortical tissues on day 5 after SW injury were immunostained with antibodies against HO-1 (red), GFAP (white), and Ror2 (Green). The lower panels are magnified views of the squares in the upper panel. Scale bars: 100 μm. (B) Neocortical tissues on day 5 after SW injury were immunostained with antibodies against HO-1 (red), GFAP (white), and Ki67 (Green). Scale bar: 100 μm.

**Figure EV3. Suppressed expression of *Ror2* reduces expression levels of multiple Nrf2-target genes in cultured astrocytes.** (A) The bar graphs show the relative expression levels of *Ftl1*, *Gclm*, *Slc7a11*, and *Pgd* in the reactive astrocytes in mouse neocortices on day 1 after MCAO or LPS injection. Data were obtained from the GEO database (GSE35338). Data are shown as the mean ± SEM (n = 4-5 animals per group). \**P*<0.05, \*\**P*<0.01, \*\*\**P*<0.001 (Tukey’s test), ns: not significant. (B) Differentiated astrocytes were stimulated with either IL-1β and TNF-α or vehicle alone in the presence or absence of bFGF for 3 days. Expression levels of *Ftl1*, *Gclm*, *Slc7a11*, and *Pgd* were analyzed by qRT-PCR. Relative expression values were determined by defining the expression levels of the indicated genes in untreated cells (-) as 1.0. The values are the mean ± SD of three independent experiments. \**P*<0.05, \*\**P*<0.01, \*\*\**P*<0.001 (Tukey’s test), ns: not significant. (C, D) Differentiated astrocytes transfected with the indicated siRNAs were stimulated with IL-1β, TNF-α and bFGF for 3 days. Expression levels of *Ftl1*, *Gclm*, *Slc7a11*, and *Pgd* were analyzed by qRT-PCR. Relative expression values were determined by defining the expression levels of the indicated genes in si-Ctrl-transfected cells as 1.0. The values are the mean ± SD of three independent experiments. \**P*<0.05, \*\**P*<0.01, \*\*\**P*<0.001 (Dunnett’s test), ns: not significant.

**Figure EV4. Basal autophagy flux is unaffected by Ror2 signaling in cultured astrocytes.** (A) Differentiated astrocytes were stimulated with IL-1β, TNF-α, and bFGF for 3 days or treated with 100 nM Bafilomycin A1 (BafA1) for 24 h. Expression levels of the indicated proteins were analyzed by Western blotting. Asterisks indicate non-specific bands. (B) Differentiated astrocytes transfected with the indicated siRNAs were stimulated with IL-1β, TNF-α, and bFGF for 2 days, then further treated with 100 nM Bafilomycin A1 (BafA1) or its vehicle (DMSO) alone for 24 h. Expression levels of NBR1 were visualized by immunostaining. Scale bar: 50 μm. (C) The bar graphs show the relative expression levels of NBR1 determined by defining those in si-Ctrl-transfected and DMSO-treated cells as 1.0. The values are the mean ± SD of 10 random fields. \*\*\**P*<0.001 (Dunnett’s test), ns: not significant. (D) Differentiated astrocytes were treated with 100 nM BafA1 or DMSO alone for 24 h. Expression levels and intracellular distribution of p62 (red) and Nrf2 (green) were visualized by immunostaining. Scale bar: 50 μm. (E) The bar graphs show the relative expression levels of p62 determined by defining those in DMSO-treated cells as 1.0. The values are the mean ± SD of 10 random fields. \*\*\**P*<0.001 (Student’s *t*-test). (F) The respective scatter dot plots show the relative intensities of nuclear Nrf2 determined by defining those in DMSO-treated cells as 1.0. The horizontal red lines indicate mean values (n > 800 cells per sample).

**Figure EV5. Damaged BBB might be repaired after day 7 of ICH.** (A) Extravasation of FITC-Dextran (green) and IgG (red) was observed around GFAP-expressing reactive astrocytes (white) in the injured striatal tissues. Asterisks indicate the injury sites. Scale bar: 100 μm. (B) The bar graphs show the relative fluorescence intensities of FITC-Dextran (left) and IgG (right) determined by defining those of contralateral sides as 1.0. Data are shown as the mean ± SD (n = 3 animals per group). \**P*<0.05, \*\*\**P*<0.001. (Student’s *t*-test). (C) Adult male mice were subjected to ICH by intrastriatal injection of collagenase. Amounts of leaked IgG heavy and light chains (IgG-H and IgG-L) in the injured (D1, 3, 5, 7, 14) or uninjured (D0) striatal tissues were analyzed by Western blotting.

